# RNA-tailor: accurate gene-level identification of transcript isoform diversity from long reads

**DOI:** 10.1101/2025.09.09.673982

**Authors:** Lilian Marchand, Hélène Touzet, Jean-Stéphane Varré

**Author notes:** Corresponding author: Lilian Marchand^1^.

## Abstract

Accurate splicing isoform identification is an essential need for progress of modern medicine and biological knowledge. The advent of long-read sequencing technologies opened up the possibility of sequencing full length transcripts. Most of the existing methods aims to predict isoforms at genome scale. However, solving all isoforms at genome scale is not always needed, mostly when one is interested in a single gene or a few genes. It also brings algorithmic constraints, encouraging the use of less accurate alignment algorithms and downgrading sensitivity of methods to determine the repertoire of isoforms. The aim of RNA-tailor is to make available an easy-to-use tool to perform single gene resolution of alternative splicing isoform repertoire with high accuracy. To leverage such precision, RNA-tailor uses a combination of exact alignment algorithm and context aware alignment corrections. By analyzing both real and simulated datasets, we show that RNA-tailor is able to achieve higher sensitivity.

## 1 INTRODUCTION

Alternative splicing (AS) is a fundamental mechanism that regulates gene expression and drives protein diversity in eukaryotes. During this process, the spliceosome selectively includes or excludes different combinations of exons from premRNAs to produce multiple mature mRNA variants from a single gene (Breitbart et al., 1987). This combinatorial selection of exons enables organisms to dramatically expand their proteome diversity without increasing genome size, playing crucial roles in development, cell differentiation, immune responses, and tissue-specific functions. In contrast, aberrant splicing patterns are implicated in numerous diseases, particularly cancer, where novel splice isoforms represent promising therapeutic targets for immunotherapy (Jiang et al., 2019).

Alternative splicing can be described in five categories. The most prevalent is exon skipping (cassette exon), where an exon is either included or excluded from the final transcript. Mutually exclusive exons involve the selection of one exon from a set of alternative ones that may be adjacent or distantly located. Intron retention occurs when an intron normally removed during splicing remains in the mature mRNA. Finally, alternative donor and acceptor sites create variations at the 5′ and 3′ splice sites respectively, often involving tandem alternative splice sites (TASS) with characteristic NAGNAG or GYNNGY sequence patterns that frequently display tissue-specific expression (Bradley et al., 2012; Wang et al., 2014; Mironov et al., 2021).

While studying splicing, three main questions are addressed. Which splicing events occur? Which isoforms are expressed? How many times each isoform is expressed? For the past two decades, RNA sequencing is mostly used to answer these questions. Several technologies are available to sequence RNA from a sample. They can be classified in two categories, short read (SR) sequencing technologies and long read (LR) sequencing technologies. While short-read (SR) RNA sequencing has dominated the field for two decades, its limited read length (typically a few hundred bases) rarely spans multiple splice junctions, necessitating computational assembly that may not accurately reconstruct full-length isoforms (Steijger et al., 2013). Long-read (LR) sequencing technologies have emerged as a transformative solution, generating reads up to 10kb that can span entire transcripts and directly reveal exon combinations. However, LR sequencing presents its own challenges: error rates can reach 10% for Oxford Nanopore Technology, complicating precise splice junction identification, and technical artifacts often produce truncated reads that fail to capture complete transcripts (Dong et al., 2023). This raises the problem of finding true isoforms, sharing a same intron chain, against those included in others. Despite these limitations, the ability to directly observe major splicing events through full-length reads represents a significant advance over short-read assembly-based approaches. Moreover, one can expect that the identification of the most abundant splicing events, such as intron retention, cassette exons, and alternative start-stop sites, can be achieved using long reads through direct observations.

The surge in long-read RNA sequencing has spawned numerous computational methods for isoform identification, including FLAIR (Tang et al., 2020), Bambu (Chen et al., 2023), IsoQuant (Prjibelski et al., 2023), ESPRESSO (Gao et al., 2023), and FREDDIE (Orabi et al., 2023). These tools share a common workflow: aligning reads to a reference genome, filtering or correcting alignments, and clustering reads to identify isoforms. Some of them also provide quantification results. The difference between these methods is their reliance on existing annotations. Annotation-guided approaches (FLAIR, Bambu, IsoQuant, ESPRESSO) generally achieve superior performance in detecting full-length transcripts (Su et al., 2024). They leverage known splice sites to correct alignment errors—for instance, FLAIR and IsoQuant adjust predicted junctions to match nearby annotated sites, Bambu employs machine learning to validate splice junctions within 10bp of known sites, and ESPRESSO reassigns low-confidence junctions within 35bp windows of high-confidence annotated junctions.

While using annotations allows to achieve better results in terms of both sensitivity and precision, this dependency on annotations becomes a liability when databases are incomplete or inaccurate, potentially increasing false positive rates. In contrast, annotation-free methods like FREDDIE rely solely on read alignments and clustering algorithms, while reference-free approaches such as isONform (Petri and Sahlin, 2023), RATTLE (Rubia et al., 2022), and CARNAC-LR (Marchet et al., 2018) perform all-against-all read clustering without genome alignment.

These various methods operate at the genome-wide level to analyze splicing dynamics from an LR RNA-seq experiment. In contrast, in this paper we focus on the problem of the identification of the repertoire of isoforms for a single gene of interest.

Here, we present RNA-Tailor, a fundamentally different approach to isoform identification that addresses key limitations of existing methods. Unlike genome-wide tools that must balance computational efficiency with accuracy across thousands of genes, RNA-Tailor focuses on the comprehensive characterization of splicing at gene level. This gene-centric approach is particularly valuable for targeted studies where researchers need to understand the complete isoform repertoire of specific genes of interest, such as disease-associated genes or therapeutic targets. This targeted strategy enables three critical advantages: First, RNA-Tailor can employ computationally intensive but highly accurate global pairwise spliced alignments that would be prohibitive at genome scale. Second, it implements correction algorithms specifically designed for long-read splice junction alignment errors. Third, RNA-Tailor operates completely annotation-free, relying exclusively on sequence similarity rather than potentially incomplete or biased reference annotations. This philosophy makes RNA-Tailor equally suited for both well-annotated model organisms and newly sequenced species lacking comprehensive annotations. By fully leveraging improvements in long-read sequencing quality and alignment algorithms, RNA-Tailor provides an unbiased, user-friendly high-resolution view of the gene complete isoform list for understanding complex splicing patterns.

## 2 MATERIALS AND METHODS

### 2.1 Algorithm

RNA-Tailor is a novel tool designed to precisely inventory the repertoire of alternatively spliced transcripts for a gene of interest from long reads. RNA-tailor takes as input an LR RNA-seq dataset and either the reference genome and the locus of the gene of interest or the reference sequence of the gene. RNA-tailor runs in three main steps, using both existing state-of-the-art alignment methods as well as homemade specific steps. It also includes refinement steps for higher accuracy. In the following, we provide details about the RNA-tailor pipeline. Figure 1 provides an overview of the key steps involved.

**Figure 1.**
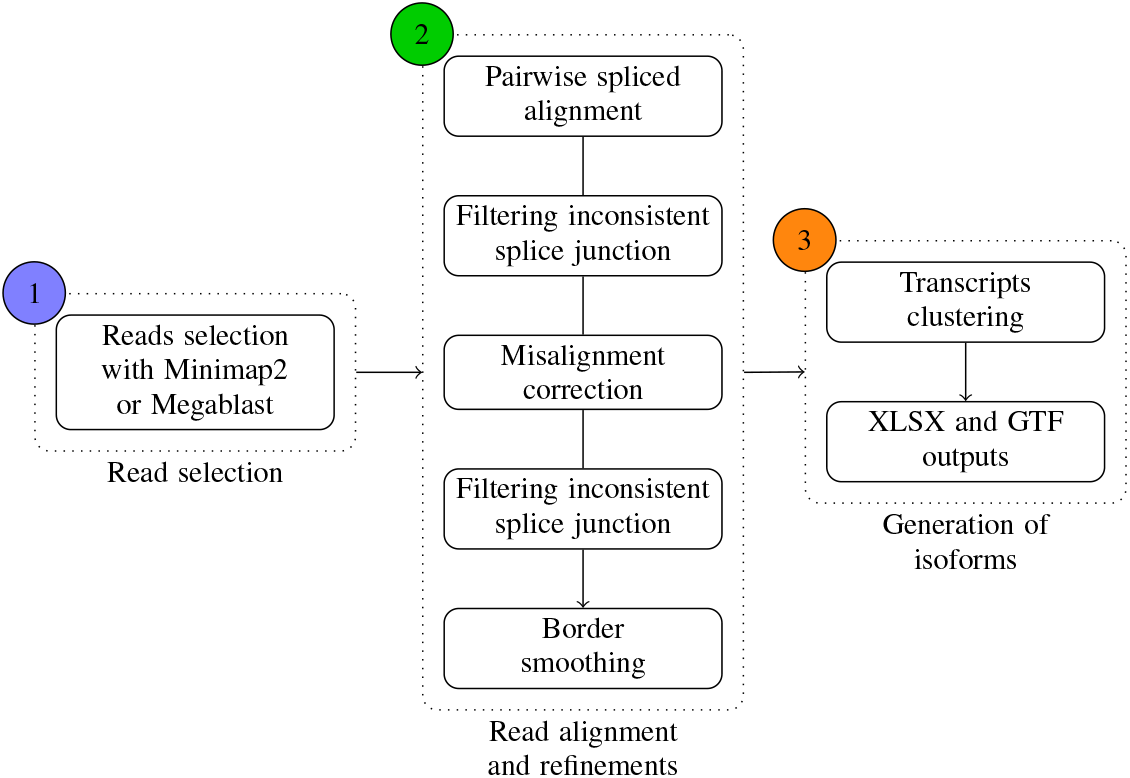
Overview of the RNA-tailor pipeline.

#### 2.1.1 Read selection

RNA-tailor provides two methods for selecting reads from the input set that correspond to the expression of the target gene. The choice depends on the type of reference sequence. If the reference sequence is the genomic sequence of the gene of interest, the use of megablast is recommended. If the reference sequence is the whole genome, the use of minimap2 is recommended. In both cases a read is selected if at least half of its bases are aligned against the genomic sequence of the target gene. Moreover a spliced alignment is performed with exonerate, and reads that do not have at least one predicted intron in their alignment (i.e. single exon reads) are discarded.

#### 2.1.2 Read alignment and refinements

From now on, we have a set of selected reads that are putative alternative transcripts of the gene of interest. We assume that reads from the same gene should share a structural homogeneity in their splicing junctions. We thus compute a preliminary prediction of splice junctions using a splice-aware alignment software, that is exonerate (using the est2genome model) (Slater and Birney, 2005). This approach sets RNA-Tailor apart from other long-read isoform identification tools, which typically recommend or directly use minimap2 for splice alignment. Unlike minimap2, which relies more on heuristics, exonerate uses an exact alignment algorithm, making it more accurate but more computationally intensive. The rationale for using exonerate stems from the fact that RNA-tailor is designed for studying of a single gene of interest, thus limiting the number of reads to be analyzed.

At this point, we have draft predicted splice junctions for each read. Indeed, spliced alignments may still contain errors, and further work is needed to improve the splice junction prediction at base resolution.

We first start by filtering reads with inconsistent splice junctions. We filter out reads with more than a quarter of isolated splice junctions (splice junctions that are more than 20bp away from the nearest junction point in other reads). Figure 2 gives an illustration. Because of sequencing errors in reads and misalignments from exonerate, we continue with correction steps of the alignments. We focus on segments of reads that could improve splice junctions support if they were realigned elsewhere. Figure 3 gives an illustration. Depending on the length of such segments, we compute a local alignment between the segment in concern and a subpart of the gene, or the segment in concern is moved to a better position. After that, we will once again filter out the reads with isolated splice junction points. Finally, as some splice junctions may not be well aligned with the reference, we apply a majority rule to decide the position of the splice site on the reference sequence (see Figure 4).

**Figure 2.**
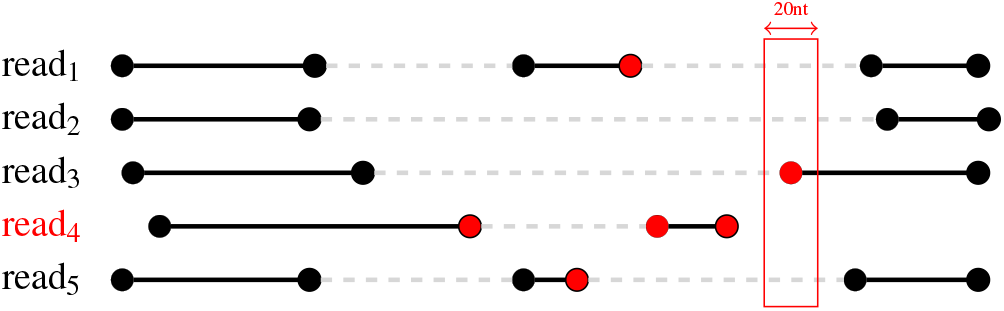
Illustration of junction point filtering. Junction points are represented by circles. The red ones are the isolated ones. Reads 1, 3 and 4 have less than 25% of isolated junction points, so they are retained. Read 2 has more than 25% of isolated junction points, so it is removed. read_1_ read_2_ read_3_ read_5_ before realignment read_1_ read_2_ read_3_ read_5_ after realignment

**Figure 3.**
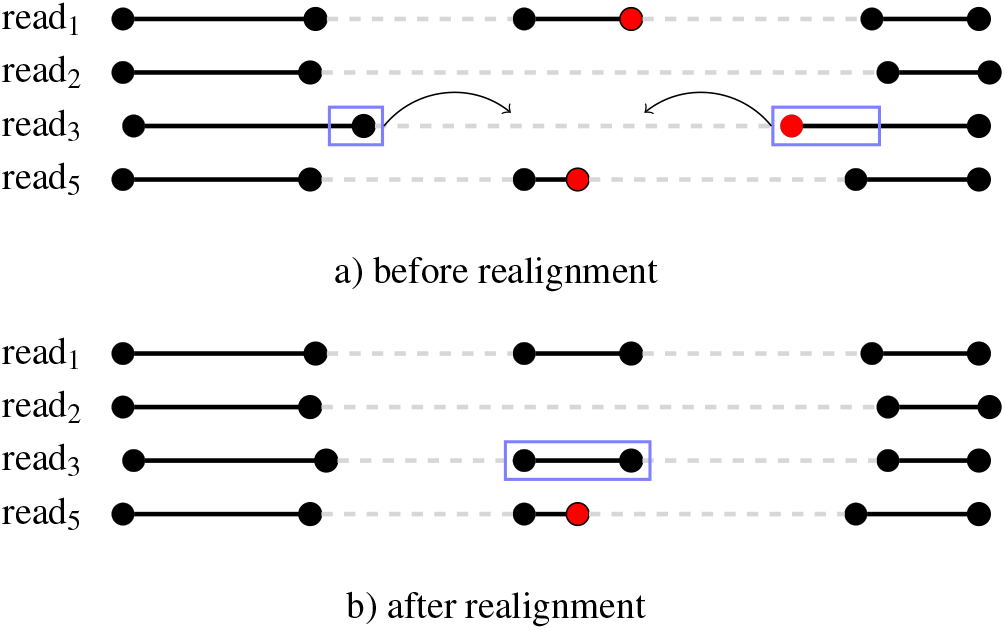
Illustration of realignment. a) Rectangles show segments of read_3_ that could be realigned. b) As long as the new alignment improves the support for splice junctions points (reducing the number of isolated junction points), it is validated. iso_1_ iso_2_ iso_3_ gene read_1_ read_2_ read_3_ read_5_

**Figure 4.**
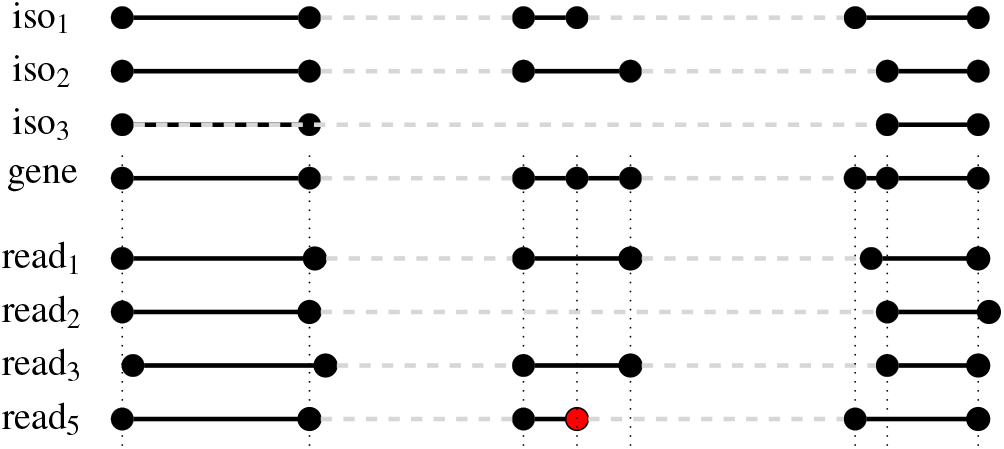
Illustration of border smoothing and determination of isoforms. Dotted lines show how border smoothing acts. Among the junction points in the same small interval, we select the majority. This gives the exonic structure of the gene. Isoforms can be determined : iso_1_ is supported by read_5_, iso_2_ is supported by read_1_ and read_3_, while iso_3_ is supported by read_2_.

#### 2.1.3 Generation of isoforms

The final step consists in the identification of isoforms. Indeed, at this step, we have a set of reads and the splice sites positions identified in the reference sequence for each of them. We have to recover isoforms from this set. We consider that two reads come from the same isoform if they share the same *intronic structure* (i.e. they share exactly the same introns, regardless of the beginning of the first exon and the end of the last exon). Figure 4 gives an illustration. An isoform is considered trustful if it is supported by at least two reads. Consequently, we output two sets of isoforms.

### 2.2 Implementation details and output

The pipeline is developped using a Snakemake framework, using Conda, Python and Biopython. It requires to have megablast and/or minimap2 installed, exonerate version 2.3 and Seqtk to be installed. All parameters can be customized via a JSON file. It outputs standard GTF files: one with isoforms supported by 1% of the total number of reads or at least 2 reads, and another one with the remaining ones. RNA-tailor provides also a XLSX output allowing to observe each read individually where blocks and junction points are given. RNA-tailor can be downloaded from https://gitlab.univ-lille.fr/bilille/RNA-tailor (scripts to reproduce the results presented in this paper are available from the https://gitlab.univ-lille.fr/RNA-tailor/RNA-tailor-publication)

### 2.3 Datasets

#### 2.3.1 Real dataset

The real dataset comes from Freddie’s publication (SRR15899612). This dataset is known to contain expressed alternative splicing isoforms. These data come from Oxford Nanopore sequencing (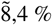 error rate) and contain splicing events that have been validated by short-read sequencing. We studied a pool of 280 genes expressed from this dataset, derived from the 294 genes selected in Freddie’s publication, excluding those with only single isoform or single exon transcripts. From these genes, we selected the known transcripts from the annotation *homo_sapiens* GRCh38 release 111 with the tags “protein coding” and “basic”, “Ensembl_canonical” or “MANE_Select”. This selection filtered out incomplete transcripts and redundancy in the isoforms of the dataset. The final dataset is made of 875 isoforms coming from 280 genes and having 868 unique intronic structures and no single exon transcript. This dataset is reffered to as “875 “ is the following. With real datasets, we do not know which isoform is actually expressed, making comparison to a ground-truth impossible. Therefore, we took advantage from the short read dataset (SRR15899613), generated from the same experiment. A custom ground-truth was created by retaining only the short reads that span at least one intron. A predicted splice junction is valid if at least one short read is aligned over it. This dataset is reffered to as “875_/SR_“ is the following. Number of genes and transcript of both datasets are given Table 1.

**Table 1.**
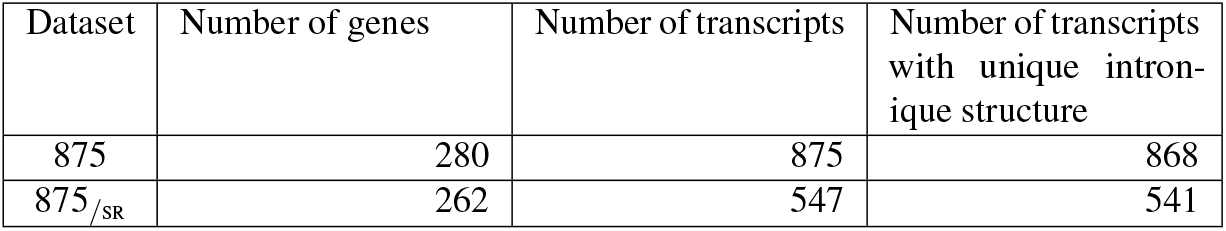
Number of genes and transcripts in reference datasets.

#### 2.3.2 Simulated dataset

The use of synthetic datasets allows to control both error rate and expression level. It also allows to simulate specific splicing events. We focused on simulating reads using the same set of 280 genes of interest following a uniform read expression profile of 20 reads per isoform. To simulate third-generation RNA sequencing reads, we used the long-read simulator PBSIM3 (Ono et al., 2022) with the ONT degradation profile “errhmm”. The transcript sequences from which the reads are sampled were extracted from the GRCh38 genome assembly. The experiments were performed with simulated reads at 5% and 10% error rate. We observed that PBSIM3 exhibits a read length distribution comparable to that of the real dataset (not shown). We also built a set of exact perfect reads, namely 0% error rate in the following, containing the known transcripts themselves (full-length) using gffread (Pertea and Pertea, 2020).

#### 2.3.3 Artificial splicing event generation

In order to evaluate the ability of the methods to detect AS events, we generated synthetic transcripts by making random splicing events in known isoforms. The three most common types of splicing modifications were chosen to simulate: Exon Skipping (EC), Intron Retention (IR), and Alternative Start/Stop (ASS). To simulate EC and IR, the number of new transcripts is randomly chosen based on the number of known transcripts for each gene: it ranges from zero to half of the number of known transcripts. Each new isoform can have only one type of modification (none contains both ASS and EC or ASS and IR, or EC and IR). EC is simulated by randomly deleting of one or more internal exons ranging from one to one third of the total number of exons. Building artificial IR involves the fusion of two consecutive exons to simulate intron retention. One exon is randomly drawn, excluding the first and last exons, and merged with the following exon. Simulating an ASS consists in searching for alternative canonical splice sites within 12bp upstream and downstream of each internal exon of each transcript. We randomly retain only one transcript per gene to simulate an ASS.

The simulated splicing events are recorded in two GTF files of modified transcripts: one for ASS and one for both EC and IR. Each file is then combined with the GTF file of known transcripts and is provided as input to PBSIM3 to simulate the reads. The number of modifications performed in our experiments is summarized Table 2.

**Table 2.** Number of genes, isoforms and reads for both real and simulated datasets.

| Dataset | Number of genes | Number of transcripts | Number of reads | Error rate |
| --- | --- | --- | --- | --- |
| SRR15899612 | 294 | | 203,844 | $\approx 8,4\%$ |
| PBSIM_0 | 280 | 875 | $875 \times 20^1$ | 0% |
| PBSIM_5 |  |  |  | 5% |
| PBSIM_10 |  |  |  | 10% |
| PBSIM_IREC_0 | 280 | 1113 | $1113 \times 20^1$ | 0% |
| PBSIM_IREC_5 |  |  |  | 5% |
| PBSIM_IREC_10 |  |  |  | 10% |
| PBSIM_TASS_0 | 280 | 1063 | $1063 \times 20^1$ | 0% |
| PBSIM_TASS_5 |  |  |  | 5% |
| PBSIM_TASS_10 |  |  |  | 10% |

<sup>1</sup>except for perfect reads, only $10\times$

### 2.4 Metrics for performance analyses

In order to evaluate the performance of RNA-tailor we compared its results against FLAIR, Bambu (unguided) and IsoQuant (unguided).

To evaluate the predicted isoforms found by each method, or by different versions of RNA-tailor, we chose to compare them using GffCompare (Pertea and Pertea, 2020). The analysis was performed discarding duplicate isoforms. While the real datasets used in some of the experiments came from Freddie’s publication, we encountered difficulties evaluating the performance of Freddie’s predictions. They did not recall any known transcripts for any of the experiments. Consequently, we were unable to use GffCompare to effectively assess its isoform prediction performance and decided to exclude Freddie from our benchmark.

#### 2.4.1 Metrics for simulated datasets

We use known transcripts from which the reads were simulated as ground-truth for GffCompare. As we know full length transcripts, we focus on the ability of the methods to find good isoforms:

**FSM (Full Splice Match)** isoforms are defined as the ones that match all junctions with a known transcript (corresponding to class code “=“, complete, exact match of intron chain, in GffCompare’s output),

**ISM (Incomplete Splice Match)** isoforms are defined as the ones that match a subset of introns from a known transcript (corresponding to class code “c”, contained in reference - intron compatible). A predicted transcript is not considered as an ISM if the known transcript has another predicted transcript being a FSM.

#### 2.4.2 Metrics for the real dataset

Among the statistics from the GffCompare output, we look at **intron level precision** and **intron level sensibility** to evaluate the ability of the tools to find good splice junctions. The intron level sensitivity quantifies the percentage of intron boundaries aligned with all known expected positions, while precision determines the ratio of intron boundaries that correctly match known positions to all predicted intron boundaries. In addition, in order to still have an assessment of the number of FSMs, we considered that a known (annotated) transcript is expressed if all its splice junctions are supported by short reads. We thus use two references: the GTF file of knwon transcripts (the one used for experiments PBSIM 0, PBSIM 5, PBSIM 10), and the GTF file of the transcripts fully supported by SRR15899613.

Finally, to assess the viability of predicted transcripts regardless of a ground-truth, we use the number of STOP codons in the internal exons of the predicted transcripts as a proxy. The underlying hypothesis is that a predicted transcript with inaccurate exon boundaries has a chance of having a stop codon in its internal exons (the first and last exons are set aside as they have a higher chance of belonging to UTR regions). Thus we compute the number of STOP codons for the three phases of the predicted transcripts and retain the minimum value. We then examine the fraction of transcripts per gene for which there is a phase without any stop codons. For reference transcripts, a fraction of one is expected (genes that do not meet this fraction of one are excluded from the analysis).

## 3 RESULTS AND DISCUSSION

### 3.1 Simulated dataset

For simulated datasets we take advantage of a ground-truth to evaluate the performance of the tools. Table 3 shows the predictions on known (annotated) isoforms and synthetic ones.

**Table 3.**
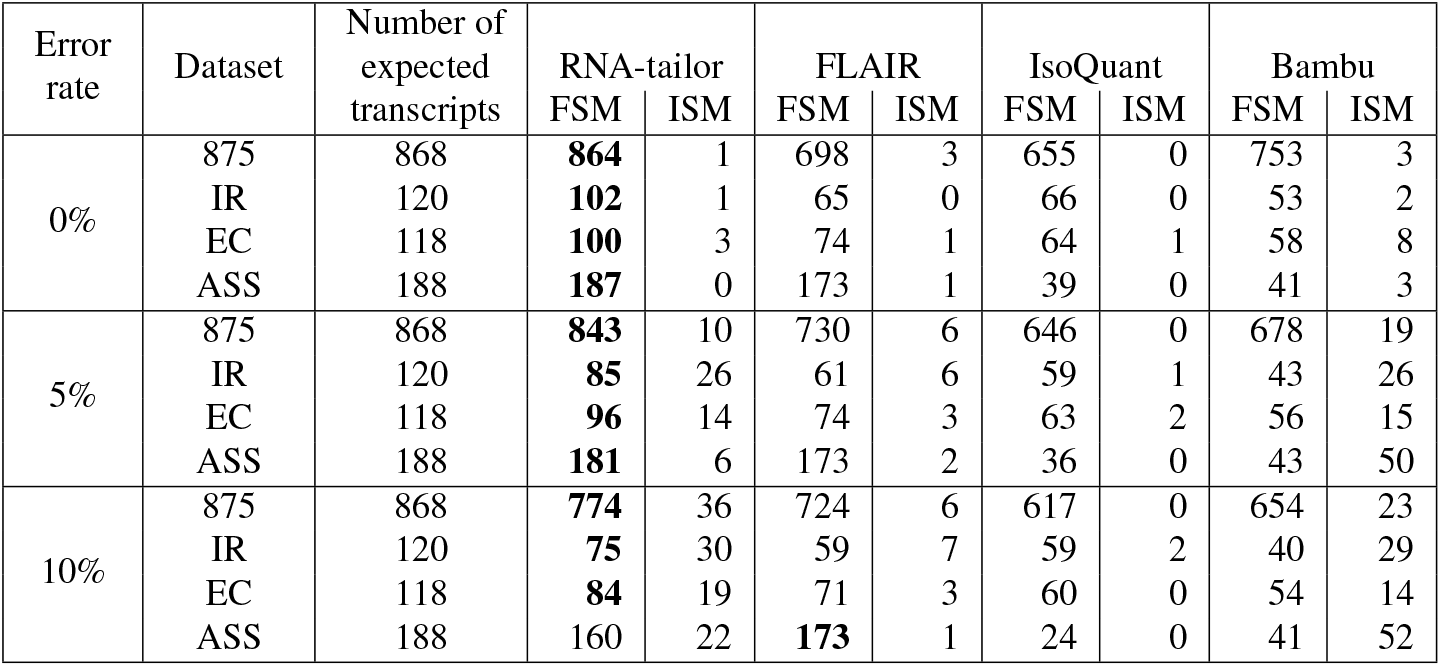
Comparison of ability to find true isoforms from RNA-tailor, FLAIR, IsoQuant and Bambu (unguided versions) on simulated datasets in terms of FSM and ISM (computed with GffCompare). Each number gives the recovered transcripts for each dataset: 875 stands for the known isoforms (datasets PBSIM (0,5,10)), IR stands for isoforms with a simulated IR (datasets PBSIM_IREC_(0,5,10)), ES stands for isoforms with a simulated ES (datasets PBSIM_IREC_(0,5,10)), and ASS stands for isoforms with a simulated ASS (datasets PBSIM_TASS_(0,5,10)) (see Table 2).

| Error rate | Dataset | Number of expected transcripts | RNA-tailor |  | FLAIR |  | IsoQuant |  | Bambu |  |
| --- | --- | --- | --- | --- | --- | --- | --- | --- | --- | --- |
|  |  |  | FSM | ISM | FSM | ISM | FSM | ISM | FSM | ISM |
| 0% | 875 | 868 | **864** | 1 | 698 | 3 | 655 | 0 | 753 | 3 |
|  | IR | 120 | **102** | 1 | 65 | 0 | 66 | 0 | 53 | 2 |
|  | EC | 118 | **100** | 3 | 74 | 1 | 64 | 1 | 58 | 8 |
|  | ASS | 188 | **187** | 0 | 173 | 1 | 39 | 0 | 41 | 3 |
| 5% | 875 | 868 | **843** | 10 | 730 | 6 | 646 | 0 | 678 | 19 |
|  | IR | 120 | **85** | 26 | 61 | 6 | 59 | 1 | 43 | 26 |
|  | EC | 118 | **96** | 14 | 74 | 3 | 63 | 2 | 56 | 15 |
|  | ASS | 188 | **181** | 6 | 173 | 2 | 36 | 0 | 43 | 50 |
| 10% | 875 | 868 | **774** | 36 | 724 | 6 | 617 | 0 | 654 | 23 |
|  | IR | 120 | **75** | 30 | 59 | 7 | 59 | 2 | 40 | 29 |
|  | EC | 118 | **84** | 19 | 71 | 3 | 60 | 0 | 54 | 14 |
|  | ASS | 188 | 160 | 22 | **173** | 1 | 24 | 0 | 41 | 52 |

#### Known isoforms identification

RNA-tailor achieves the best number of FSM recoveries with a sensitivity between 89% (10% error rate) and 99% (with 0% error rate). Overall, FLAIR performs better than IsoQuant and Bambu as soon as reads contain errors. Interestingly, even when we provide exact full-length transcripts (0% error rate in reads), none of the methods are able to recover all the isoforms. This underscores the difficulty of accurately capturing the signal.

#### Artificial IR and EC identification

For the identification of artificial IR and EC events, RNA-tailor consistently outperforms other tools. This greater sensitivity of RNA-tailor can be attributed to its ability to extract, correct, and preserve the signal provided by exonerate.

#### Artificial ASS identification

Detection of ASS events is certainly the most challenging task for alternative isoform identification tools, as such events involve very small regions (a few bases) around splice junctions. Deciphering the signal from noisy data is hard. Sequencing and alignment errors occurring in those regions have a critical impact on the detection of the good splice junction and consequently on the discovery of alternative isoforms. RNA-tailor identifies roughly as many ASS events as FLAIR unguided in each experiment, while IsoQuant and Bambu perform the worst.

#### Incomplete Splice Matches

The number of ISMs provides a good approximation of the number of missed predictions. The number of ISMs if very low for FLAIR and IsoQuant while it is higher for RNA-tailor and Bambu, suggesting that RNA-tailor and Bambu have a higher sensitivity. Indeed, the ISMs can be seen as a way to identify methods that can detect signals from new alternative events. RNA-tailor is the only one for which the addition of the number of FSMs and ISMs almost reaches the maximum number of expected predictions.

#### Does read correction improve predictions ?

When working with LR, the alignment quality is impacted by the high sequencing errors. Sequence correction generally improves the quality of the spliced alignment of the reads to the reference gene, increasing the support for splice junctions. We thus tested this hypothesis by applying auto-correction to the selected reads using isONcorrect (Sahlin et al., 2021) (between step 1 and 2, Figure 1). Resultats are given Supplementary table 6. Overall, applying read correction with RNA-tailor on knwon transcripts or simulated ones with IR and ES events tends to remove FSMs while the sum of FSMs and ISMs remains the same. Overall, the analysis of prediction results for these two types of events highlights RNA-tailor’s ability, supported by exonerate’s sensitivity, to distinguish genuine signals from noise, a key capability for accurate and sensitive RNA identification. Applying read correction on simulated ASS datasets removes about 30% of the number of FSMs, and about 7% of the sum of FSMs and ISMs. This could be expected as preserving ASS while applying read correction is challenging due to their small size and direct impact on splice site selection.

#### Comparison to guided versions

FLAIR, IsoQuant and Bambu can take as input an annotation file to help predictions. Results of the running of these tools in their guided version compared to RNA-tailor is given as Supplementary Table 7. Considering the predictions of the set known transcripts, it is noteworthy that without prior knowledge RNA-tailor achieves better results than FLAIR, and as good results as IsoQuant and Bambu (except when the read error rate is 10%). Considering the predictions of new transcripts with simulated events, all the tools performs poorly compared to RNA-tailor. This is even more true for ASS events for which there is a complete lack of detection of novel ASS sites.

### 3.2 Real dataset

We recall that for the real dataset we do not know which transcript is expressed. We thus use boths dataset 875 and dataset 875_/SR_ as reference, the last considered as a ground-truth. Table 4 gives results of predicted isoforms from Bambu, FLAIR, Freddie, IsoQuant and RNA-tailor. RNA-tailor and FLAIR are the only ones predicting isoforms for each gene.

**Table 4.** Results of predictions for each tool. A predicted transcript is associated to gene if its bounds overlap gene bounds (top compared to dataset 875, bottom compared to dataset 875_/SR_).

|  | Bambu | FLAIR | Freddie | IsoQuant | RNA-tailor |
| --- | --- | --- | --- | --- | --- |
| Number of predicted transcripts | 851 | 8273 | 2923 | 589 | 4122 |
| Dataset 875 (280 genes) |  |  |  |  |  |
| Number of predicted transcripts associated to a gene | 632 | 6270 | 1559 | 506 | 4121 |
| Ratio | 74% | 76% | 53% | 86% | **100%** |
| Number of genes having at least an associated transcript | 245 | 280 | 187 | 218 | 280 |
| Ratio | 88% | **100%** | 67% | 78% | **100%** |
| Dataset 875<sub>/SR</sub> (262 genes) |  |  |  |  |  |
| Number of predicted transcripts associated to a gene | 605 | 5689 | 1438 | 498 | 3836 |
| Ratio | 71% | 69% | 49% | 85% | **93%** |
| Number of genes having at least an associated transcript | 235 | 262 | 174 | 210 | 262 |
| Ratio | 90% | **100%** | 66% | 80% | **100%** |

#### Ability to find good splice junctions

We count the number of times intron bounds from a reference file are retrieved to estimate the ability of a tool to capture good splice junctions (Figures 5a and 5b). RNA-tailor achieves the best ratio between sensitivity and precision while FLAIR has a very low precision, and IsoQuant and Bambu have a low sensitivity.

**Figure 5.**
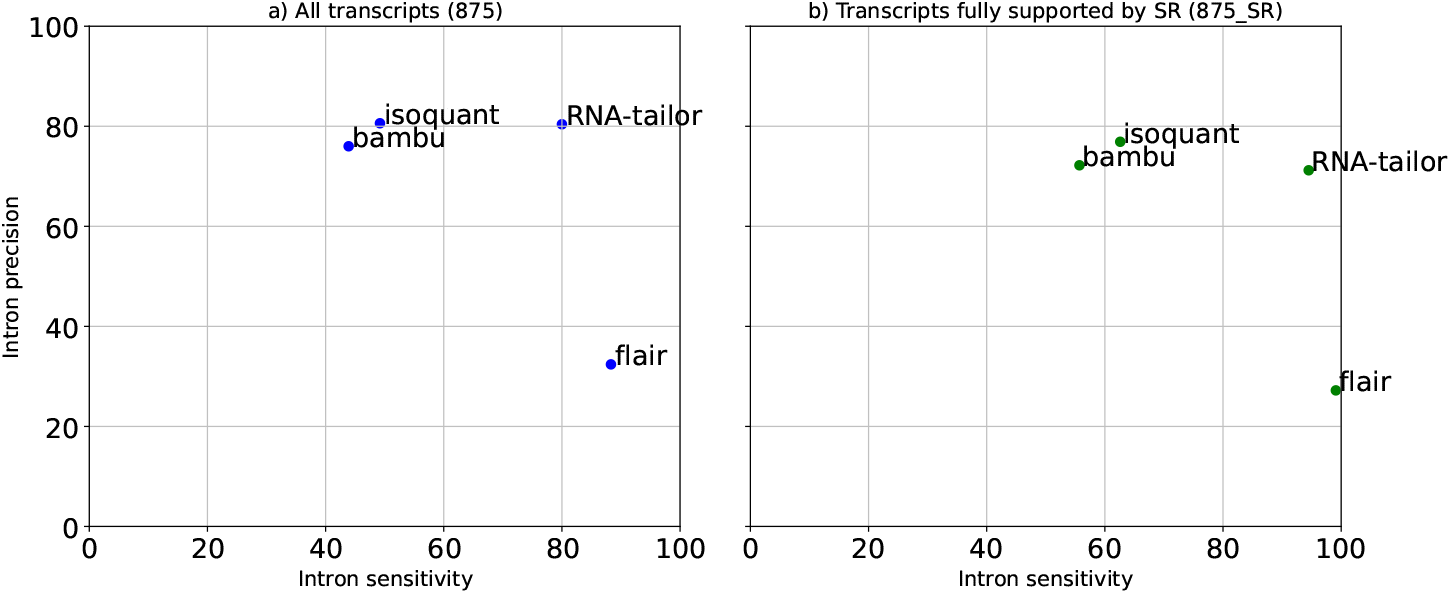
Intron and exon sensitivity and precision recovery for RNA-tailor, FLAIR, IsoQuant and Bambu. a) on the dataset SRR15899612 with 875 as reference, b) on the dataset SRR15899612 with 875_/SR_ as reference.

#### Ability to find good isoforms

Since we cannot compare the predicted isoforms with a ground-truth, we wanted to test if the predicted isoforms have a phase in which there is no STOP codon in internal exons (see Methods above). Table 5 gives statistics about the number of putative CDS spanning all internal exons. RNA-tailor achieves clearly better performance considering the number of genes for which such transcripts are predicted, while Bambu achieves a slighly better performance than RNA-tailor considering the number of transcripts. This can be explained by the higher precision of Bambu and the higher sensitivity of RNA-tailor. It is thus interesting to note than, despite the high number of predicted trancripts by RNA-tailor (more than 3000), half of them dos not contain STOP codon in their internal exons, which is not the case for FLAIR, another method with good sensitivity. Finally, figure 6 reports the fraction of predicted isoforms having 0 STOP codon in internal exons for each gene for each tool. We observe that FLAIR and Freddie tend to produce transcripts that does not meet the non codon stop criteria in internal exons, which confirms their lack of precision. The results for Freddie highlight a problem with the accuracy of splice junction predictions, as expected from the null FSM counts on simulated data. IsoQuant achieves good results for some genes and bad results for others. The trend remains the same as described above for Bambu and RNA-tailor, the difference comming from the different balance between sensitivity and precision.

**Table 5.** Analysis of transcripts associated to a gene in reference datatsets having at least 3 exons (top compared to dataset 875, bottom compared to dataset 875_/SR_). The number of predicted transcripts without a STOP codon gives the number of transcript for which there is a phase such that no stop codon are found in internal exons.

|  | Bambu | FLAIR | Freddie | IsoQuant | RNA-tailor |
| --- | --- | --- | --- | --- | --- |
| Dataset 875 (280 genes) |  |  |  |  |  |
| Number of predicted transcripts associated to a gene | 526 | 4738 | 1141 | 498 | 3693 |
| Number of genes having at least an associated transcript | 223 | 266 | 178 | 217 | **267** |
| Number of predicted transcripts without a STOP codon | 298 | 1297 | 200 | 219 | 1955 |
| Ratio | **57%** | 27% | 18% | 44% | 53% |
| Number of genes having at least an associated transcript without a STOP codon | 151 | 228 | 107 | 119 | 258 |
| Ratio | 68% | 86% | 60% | 55% | **97%** |
| Dataset 875<sub>/SR</sub> (262 genes) |  |  |  |  |  |
| Number of predicted transcripts associated to a gene | 508 | 4304 | 1054 | 490 | 3433 |
| Number of genes having at least an associated transcript | 215 | 259 | 165 | 209 | **250** |
| Number of predicted transcripts without a STOP codon | 292 | 1206 | 188 | 215 | 1808 |
| Ratio | **57%** | 28% | 18% | 44% | 53% |
| Number of genes having at least an associated transcript without a STOP codon | 146 | 212 | 101 | 115 | **241** |
| Ratio | 68% | 82% | 61% | 55% | **96%** |

**Figure 6.**
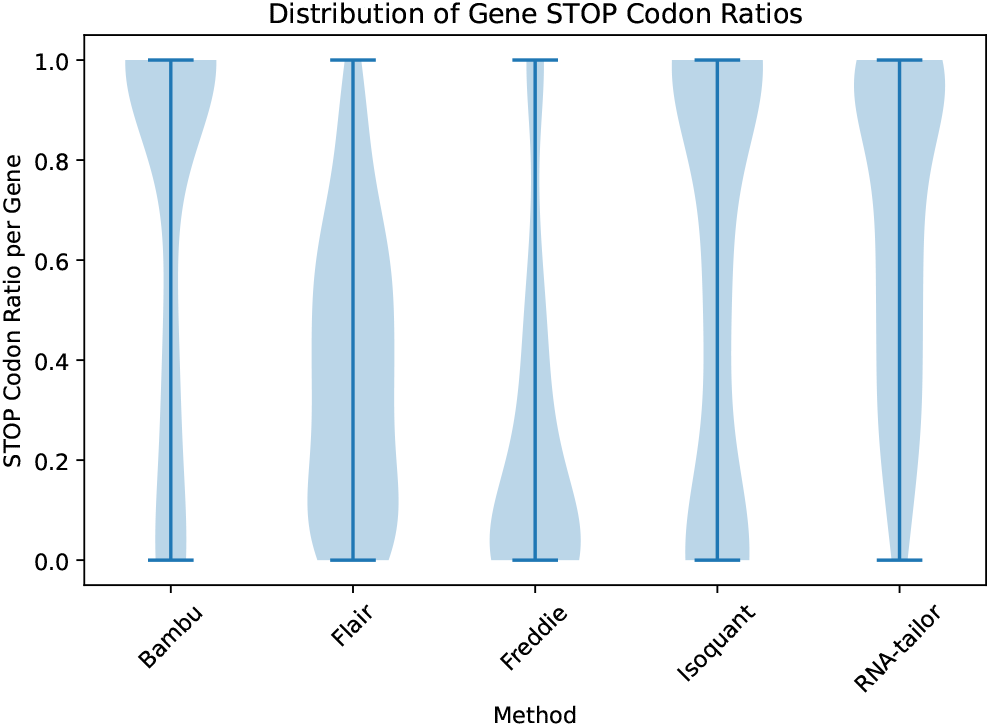
Fraction of predicted isoforms for which there is a phase with 0 STOP codon in internal exons for each tool. Only isoforms associated to a gene of dataset 875_/SR_ with at least 3 exons were analysed.

#### Comparison to known annotations

To end the analysis on the real dataset we compared the predicted isoforms in terms of FSMs and ISMs with annotations from 875 and 875_/SR_. The results presented should thus be compared between tools, and not interpreted as the ability of a method to find the truly expressed isoforms. We computed the number of FSMs and ISMs in comparison with both on all known isoforms and on knwon isformorms whose all splice junctions are fully supported by short read splice junction alignments. The results of this experiment are presented Figure 7. RNA-tailor has the highest number of FSMs and ISMs whatever the reference used. This demonstrates the ability of RNA-tailor to discover alternative isoforms without prior knowledge.

**Figure 7.**
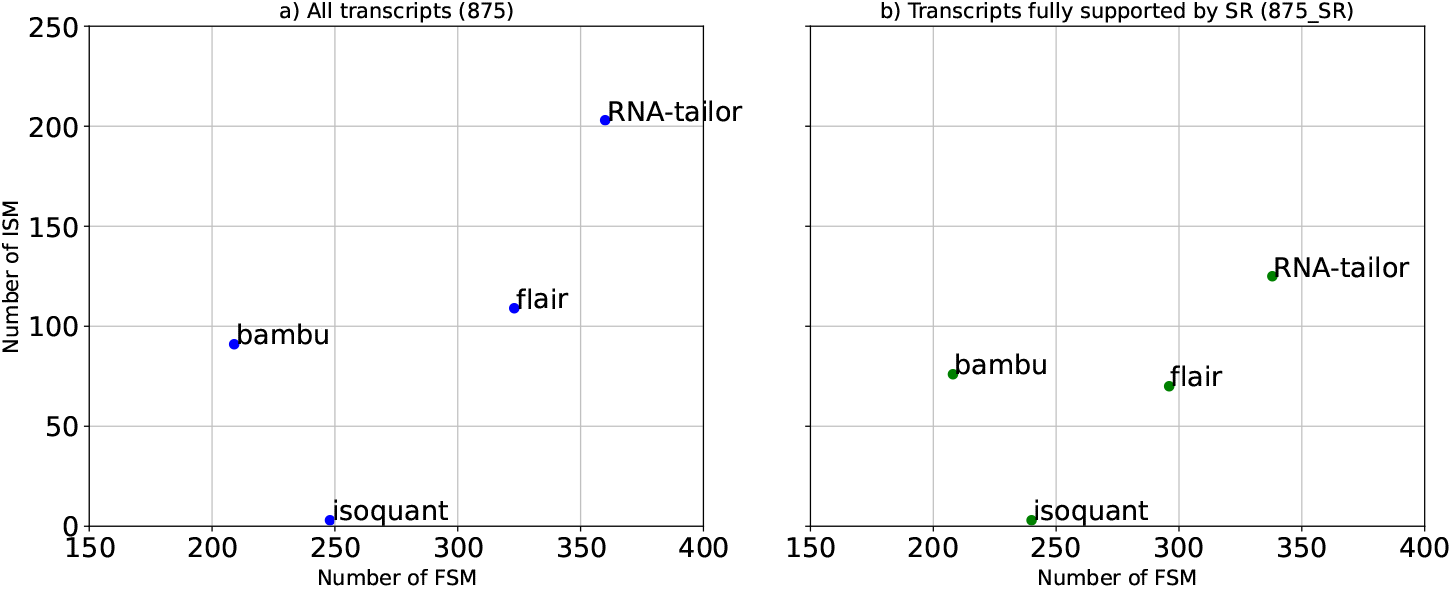
Number of FSMs and ISMs recovered by RNA-tailor, FLAIR, IsoQuant and Bambu. a) on the dataset SRR15899612 with 875 as reference, b) on the dataset SRR15899612 with 875_/SR_ as reference.

### 3.3 Use case

While previous sections analyzed a high number of genes, the aim of RNA-tailor is to precisely analyse a given gene. We show two experiments. Firstly, we ran RNA-tailor with a set of reads coming from mouse (dataset) against a reference gene coming from mouse (ENSMUSG00000000827). Secondly we ran RNA-tailor the same set of reads but against a reference gene coming from rat (ENSRNOG00000080504). The resulting alignment and splice junction predictions are depicted Figure 8. For the first experiment, initially, 26 reads were selected, 11 reads remained after filtering inconsistent splice junctions and 9 reads remained after border smoothing. Realignment has been done for read 477. For the second experiment, initially, 32 reads were selected, 8 reads remained after filtering inconsistent splice junctions and 6 reads remained after border smoothing. Realignment has been done for read 82. All the splice junctions (intron boudaries) correspond exactly to annotated splice junctions for both experiments. The mouse vs. mouse experiment demonstrates the ability of RNA-tailor to find accurately splice junctions without prior knowledge. The comparison of results from mouse vs. rat against mouse vs. mouse demonstrates the ability of RNA-tailor to inventory a repertoire of transcripts even in the case only a genomic sequence of a close species is available.

**Figure 8.**
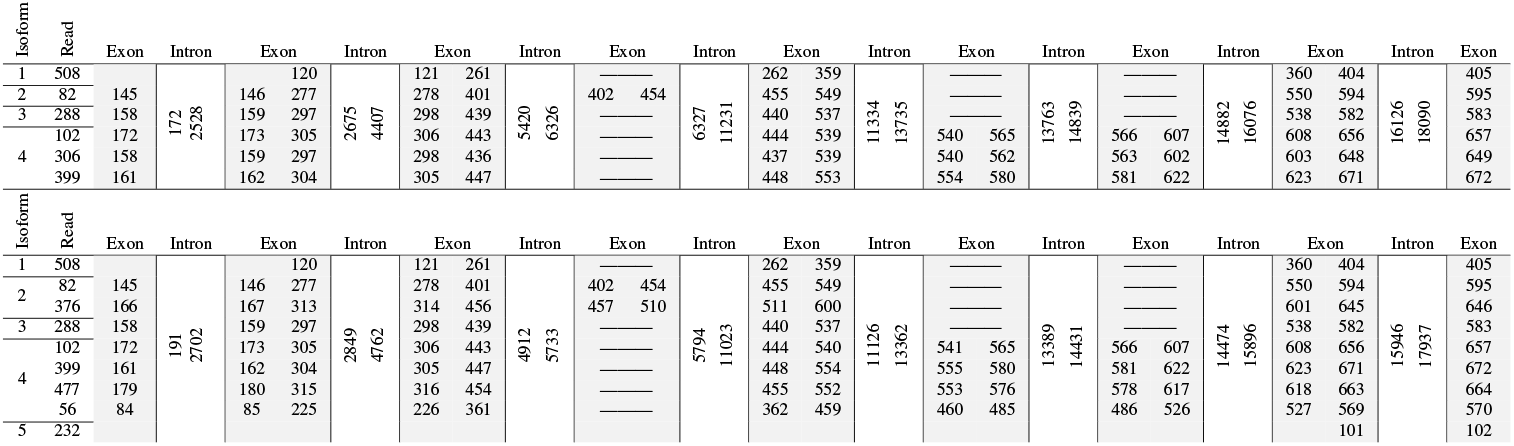
Predicted isoforms found by RNA-tailor with mouse long reads against rat genomic sequence ENSRNOG00000080504 (top) and mouse genomic sequence ENSMUSG00000000827 (bottom). Exon boudaries are given relatively to the begining of the reads. Intron boundaries correspond to splice junctions predicted by RNA-tailor, positions are given relatively to the begining of the genomic sequence. Spreadsheet files produced by RNA-tailor are given as supplementary files.

## CONCLUSION

In this work, we introduced RNA-tailor, a targeted, annotation-free approach for transcript isoform identification at the gene level. RNA-tailor represents a tool of choice when researchers are specifically interested in characterizing the complete isoform diversity of individual genes rather than conducting genome-wide analyses. Through extensive benchmarks on both simulated and real datasets, we demonstrated that RNA-tailor achieves a superior balance of sensitivity and precision compared to existing methods, including FLAIR, Bambu, IsoQuant, and Freddie. Its ability to recover both known and novel isoforms, even under high error rates, highlights the robustness of the underlying strategy, which combines exact spliced alignments with context-aware refinement steps. Importantly, RNA-tailor ‘s independence from reference annotations allows accurate isoform discovery in both well-annotated model organisms and species with limited genomic resources.

Beyond outperforming competing tools, RNA-tailor also proved versatile in practical applications. In our use case, it successfully reconstructed isoforms using a rat genomic sequence as reference for mouse reads, demonstrating that the tool can leverage sequences from closely related species as effective alignment tutors. This capability expands the scope of transcriptomic analyses to organisms with incomplete or underrepresented annotations, enabling accurate splicing studies in non-model or newly sequenced species. By creatively exploiting cross-species similarity, RNA-tailor opens the door to exploring transcript diversity in contexts where conventional annotation-guided methods remain limited.

Altogether, these results establish RNA-tailor as a reliable and versatile framework for high-resolution isoform discovery. By enabling precise reconstruction of transcript diversity, RNA-tailor provides a valuable tool to advance our understanding of alternative splicing across diverse biological contexts.

## ACKNOWLEDGMENTS

I would like to thank the ANR ASTER project for providing data, expertise, and funding. I am also very grateful to Cyprien Borée and Aymeric Antoine-Lorquin for their valuable contribution to the coding and the structural basis of the project.

## SUPPLEMENTAL MATERIAL

**Table 6.** Comparison of the number of predicted isoforms with RNA-tailor using or not isONcorrect. Results for datasets with 0% error rate are not shown since isONcorrect has no effect.

| Error rate | Dataset | Number of expected transcripts | RNA-tailor with isONcorrect |  | RNA-tailor w/ isONcorrect |  | FSM ratio |
| --- | --- | --- | --- | --- | --- | --- | --- |
|  |  |  | FSM | ISM | FSM | ISM |  |
| 5% | 875 | 868 | 811 | 18 | 843 | 10 | 96% |
|  | IR | 120 | 76 | 34 | 85 | 26 | 89% |
|  | EC | 118 | 93 | 14 | 96 | 14 | 97% |
|  | ASS | 188 | 126 | 47 | 181 | 6 | 70% |
| 10% | 875 | 868 | 747 | 38 | 774 | 36 | 97% |
|  | IR | 120 | 69 | 39 | 75 | 30 | 92% |
|  | EC | 118 | 83 | 20 | 84 | 19 | 99% |
|  | ASS | 188 | 104 | 65 | 160 | 22 | 65% |

**Table 7.**
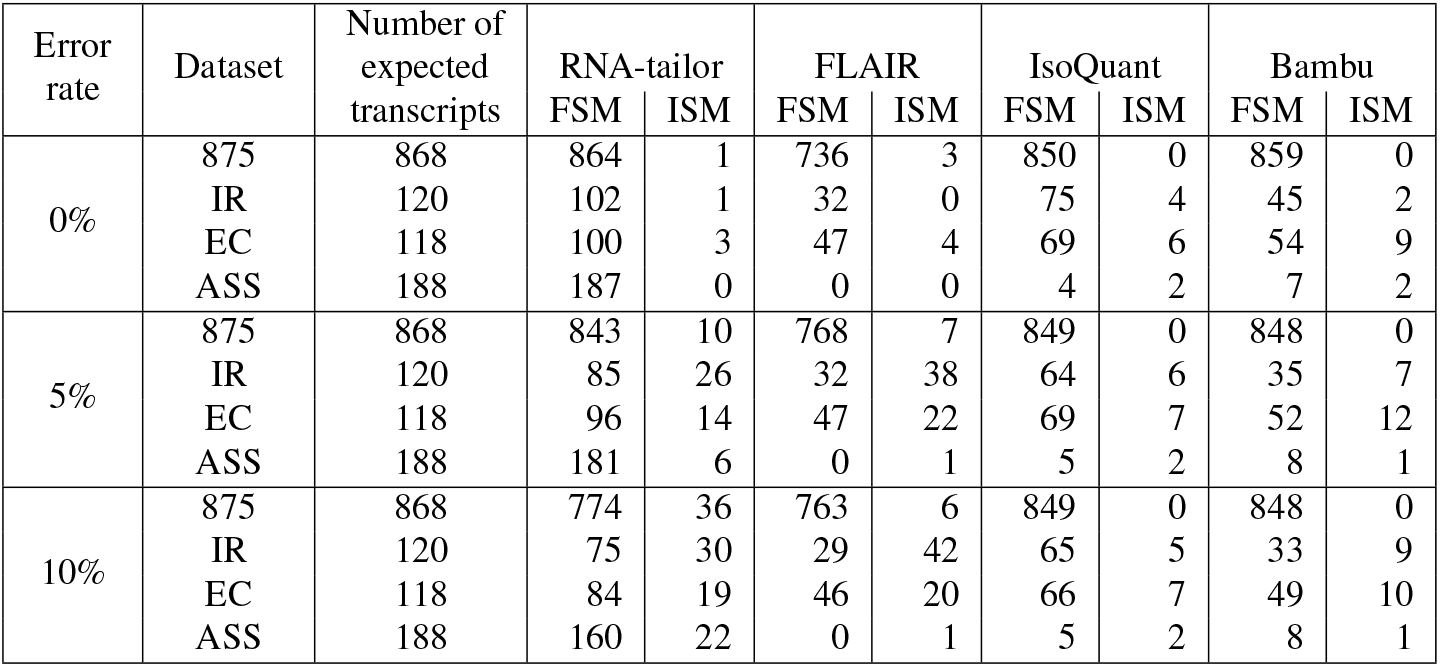
Comparison of ability to find true isoforms from FLAIR, IsoQuant and Bambu in their **guided** versions and RNA-tailor, on simulated datasets in terms of FSM and ISM (computed with GffCompare). Each number gives the recovered transcripts for each dataset: 875 stands for the known isoforms (datasets PBSIM (0,5,10)), IR stands for isoforms with a simulated IR (datasets PBSIM_IREC_(0,5,10)), ES stands for isoforms with a simulated ES (datasets PBSIM_IREC_(0,5,10)), and ASS stands for isoforms with a simulated ASS (datasets PBSIM_TASS_(0,5,10)) (see Table 2).

| Error rate | Dataset | Number of expected transcripts | RNA-tailor |  | FLAIR |  | IsoQuant |  | Bambu |  |
| --- | --- | --- | --- | --- | --- | --- | --- | --- | --- | --- |
|  |  |  | FSM | ISM | FSM | ISM | FSM | ISM | FSM | ISM |
| 0% | 875 | 868 | 864 | 1 | 736 | 3 | 850 | 0 | 859 | 0 |
|  | IR | 120 | 102 | 1 | 32 | 0 | 75 | 4 | 45 | 2 |
|  | EC | 118 | 100 | 3 | 47 | 4 | 69 | 6 | 54 | 9 |
|  | ASS | 188 | 187 | 0 | 0 | 0 | 4 | 2 | 7 | 2 |
| 5% | 875 | 868 | 843 | 10 | 768 | 7 | 849 | 0 | 848 | 0 |
|  | IR | 120 | 85 | 26 | 32 | 38 | 64 | 6 | 35 | 7 |
|  | EC | 118 | 96 | 14 | 47 | 22 | 69 | 7 | 52 | 12 |
|  | ASS | 188 | 181 | 6 | 0 | 1 | 5 | 2 | 8 | 1 |
| 10% | 875 | 868 | 774 | 36 | 763 | 6 | 849 | 0 | 848 | 0 |
|  | IR | 120 | 75 | 30 | 29 | 42 | 65 | 5 | 33 | 9 |
|  | EC | 118 | 84 | 19 | 46 | 20 | 66 | 7 | 49 | 10 |
|  | ASS | 188 | 160 | 22 | 0 | 1 | 5 | 2 | 8 | 1 |

## REFERENCES

Bradley, R. K., Merkin, J., Lambert, N. J., and Burge, C. B. (2012). Alternative Splicing of RNA Triplets Is Often Regulated and Accelerates Proteome Evolution. PLoS Biology, 10(1):e1001229.

Breitbart, R. E., Andreadis, A., and Nadal-Ginard, B. (1987). Alternative Splicing: A Ubiquitous Mechanism for the Generation of Multiple Protein Isoforms from Single Genes. Annual Review of Biochemistry, 56(1):467–495.

Chen, Y., Sim, A., Wan, Y. K., Yeo, K., Lee, J. J. X., Ling, M. H., Love, M. I., and Göke, J. (2023). Context-aware transcript quantification from long-read RNA-seq data with Bambu. Nature Methods, 20(8):1187–1195.

Dong, X., Du, M. R. M., Gouil, Q., Tian, L., Jabbari, J. S., Bowden, R., Baldoni, P. L., Chen, Y., Smyth, G. K., Amarasinghe, S. L., Law, C. W., and Ritchie, M. E. (2023). Benchmarking long-read RNA-sequencing analysis tools using in silico mixtures. Nature Methods, 20(11):1810–1821.

Gao, Y., Wang, F., Wang, R., Kutschera, E., Xu, Y., Xie, S., Wang, Y., Kadash-Edmondson, K. E., Lin, L., and Xing, Y. (2023). ESPRESSO: Robust discovery and quantification of transcript isoforms from error-prone long-read RNA-seq data. Science Advances, 9(3):eabq5072.

Jiang, T., Shi, T., Zhang, H., Hu, J., Song, Y., Wei, J., Ren, S., and Zhou, C. (2019). Tumor neoantigens: from basic research to clinical applications. Journal of Hematology & Oncology, 12(1):93.

Marchet, C., Lecompte, L., Da Silva, C., Cruaud, C., Aury, J.-M., Nicolas, J., and Peterlongo, P. (2018). Carnac-lr: Clustering coefficient-based acquisition of rna communities in long reads. In JOBIM 2018-Journées Ouvertes Biologie, Informatique et Mathématiques, pages 1–3.

Mironov, A., Denisov, S., Gress, A., Kalinina, O. V., and Pervouchine, D. D. (2021). An extended catalogue of tandem alternative splice sites in human tissue transcriptomes. PLoS Computational Biology, 17(4):e1008329.

Ono, Y., Hamada, M., and Asai, K. (2022). PBSIM3: a simulator for all types of PacBio and ONT long reads. NAR Genomics and Bioinformatics, 4(4):qac092.

Orabi, B., Xie, N., McConeghy, B., Dong, X., Chauve, C., and Hach, F. (2023). Freddie: annotation-independent detection and discovery of transcriptomic alternative splicing isoforms using long-read sequencing. Nucleic Acids Research, 51(2):e11.

Pertea, G. and Pertea, M. (2020). GFF Utilities: GffRead and GffCompare. F1000Research, 9:ISCB Comm J–304.

Petri, A. J. and Sahlin, K. (2023). isONform: reference-free transcriptome reconstruction from Oxford Nanopore data. Bioinformatics, 39(Supplement 1):i222–i231.

Prjibelski, A. D., Mikheenko, A., Joglekar, A., Smetanin, A., Jarroux, J., Lapidus, A. L., and Tilgner, H. U. (2023). Accurate isoform discovery with IsoQuant using long reads. Nature Biotechnology, 41(7):915–918. isoquant.

Rubia, I. d. l., Srivastava, A., Xue, W., Indi, J. A., Carbonell-Sala, S., Lagarde, J., Albá, M.M., and Eyras, E. (2022). RATTLE: reference-free reconstruction and quantification of transcriptomes from Nanopore sequencing. Genome Biology, 23(1):153.

Sahlin, K., Sipos, B., James, P. L., and Medvedev, P. (2021). Error correction enables use of Oxford Nanopore technology for reference-free transcriptome analysis. Nature Communications, 12(1):2.

Slater, G. S. C. and Birney, E. (2005). Automated generation of heuristics for biological sequence comparison. BMC Bioinformatics, 6(1):31.

Steijger, T., Abril, J. F., Engström, P. G., Kokocinski, F., Hubbard, T. J., Guigó, R., Harrow, J., and Bertone, P. (2013). Assessment of transcript reconstruction methods for RNA-seq. Nature methods, 10(12):1177 – 1184.

Su, Y., Yu, Z., Jin, S., Ai, Z., Yuan, R., Chen, X., Xue, Z., Guo, Y., Chen, D., Liang, H., Liu, Z., and Liu, W. (2024). Comprehensive assessment of mRNA isoform detection methods for long-read sequencing data. Nature Communications, 15(1):3972.

Tang, A. D., Soulette, C. M., van Baren, M. J., Hart, K., Hrabeta-Robinson, E., Wu, C. J., and Brooks, A. N. (2020). Full-length transcript characterization of SF3B1 mutation in chronic lymphocytic leukemia reveals downregulation of retained introns. Nature Communications, 11(1):1438.

Wang, M., Zhang, P., Shu, Y., Yuan, F., Zhang, Y., Zhou, Y., Jiang, M., Zhu, Y., Hu, L., Kong, X., and Zhang, Z. (2014). Alternative splicing at GYNNGY 5′ splice sites: more noise, less regulation. Nucleic Acids Research, 42(22):13969–13980.

